# Production of OSU G5P[7] Porcine Rotavirus Expressing a Fluorescent Reporter by Reverse Genetics

**DOI:** 10.1101/2024.01.26.577454

**Authors:** Anthony J. Snyder, Chantal A. Agbemabiese, John T. Patton

**Affiliations:** Department of Biology, Indiana University, Bloomington, IN 47405, USA; Department of Electron Microscopy and Histopathology, Noguchi Memorial Institute for Medical Research, College of Health Sciences, University of Ghana, Accra 00233, Ghana

**Keywords:** rotavirus, reverse genetics, expression vector, porcine rotavirus

## Abstract

Rotaviruses are a significant cause of severe, potentially life-threatening gastroenteritis in the young of many economically important animals. Although vaccines against porcine rotavirus exist, both live oral and inactivated, their effectiveness in preventing gastroenteritis is less than ideal. Thus, a need remains for the development of new generations of porcine rotavirus vaccines. The Ohio State University (OSU) rotavirus strain represents a *Rotavirus A* species with a G5P[7] genotype, the genotype most frequently associated with rotavirus disease in piglets. Using complete genome sequences that were determined by Nanopore sequencing, we developed a robust reverse genetics system enabling recovery of recombinant (r)OSU rotavirus. Although rOSU grew to high titers (∼10^7^ plaque-forming units/ml), its growth kinetics were modestly decreased in comparison to laboratory-adapted OSU virus. The reverse genetics system was used to generate rOSU rotavirus that served as an expression vector for foreign protein. Specifically, by engineering a fused NSP3-2A-UnaG open reading frame into the segment 7 RNA, we produced genetically stable rOSU that expressed the fluorescent UnaG protein as a functional separate product. Together, these findings raise the possibility of producing improved live oral porcine rotavirus vaccines through reverse genetics-based modification or combination porcine rotavirus vaccines that can express neutralizing antigens of other porcine enteric diseases.

## INTRODUCTION

Reverse genetics systems have been developed for several *Rotavirus A* strains, including those that infect non-human primates (SA11 and RRV), humans (KU, CDC-9, Odelia, and RIX4414-like), cattle (UK and RF), mice (rD6/2-2g), and birds (PO-13) (1–9). These systems can be used for producing next-generation live oral vaccines, dual vaccine platforms that express foreign proteins, and diagnostic tools. Rotaviruses (RV) account for a significant disease burden within important livestock, highlighting the need for effective animal rotavirus vaccines. Nonetheless, no reverse genetics systems currently exist for any porcine rotaviruses, including those with the G5P[7] genotype, which represents the most frequent cause of disease in porcine populations (10, 11).

RV is a major cause of acute gastroenteritis in piglets (10, 11). The virus is transmitted by the fecal-oral route and damages small intestinal enterocytes; as such, milk consumed by nursing piglets is not digested or absorbed into the intestines (12). Moreover, RV is resistant to environmental factors, such as temperature, pH, and common disinfectants, which creates a persistent risk of infection (13, 14). Although RV-induced diarrhea is associated with low mortality and high morbidity, productivity losses create a significant economic burden on the global pork industry. Current vaccines only control diarrhea among infected populations (10, 11). Thus, a need exists for vaccines that are more effective.

Through modification of the RV genome by reverse genetics, the virus can be used as an expression vector of foreign proteins (2, 3, 15–20). The RV genome is composed of 11 segments of double-stranded (ds)RNA. Each segment contains the coding sequence for a single protein except for segment 11, which expresses two proteins (21). As one approach for using RV as an expression vector, the NSP3 open reading frame (ORF) in the segment 7 RNA is replaced with a modified ORF that encodes NSP3 fused to a foreign protein. Insertion of the 2A translational stop-restart element allows for the modified segment 7 ORF to separately express NSP3 and the foreign protein (16–18, 22–24). This approach has been used to generate recombinant RV that express fluorescent reporters, such as UnaG [green], mRuby [red], and TagBFP [blue], and other viral proteins, such as the norovirus VP1 capsid protein and respiratory syndrome coronavirus 2 S1 spike domain (3, 15–18). Such recombinant viruses may be used to generate combination vaccines capable of inducing protective immune responses against RV and a second pathogenic virus.

We report a robust reverse genetics system for The Ohio State University (OSU) G5P[7] porcine RV. The complete sequences of its 11 dsRNA genome segments were determined by Nanopore sequencing of a laboratory-adapted OSU strain (25–27). Using T7 expression plasmids that contain coding sequences for each genome segment, we generated recombinant OSU (rOSU) and 11 rSA11/OSU monoreassortants. Single step growth curves, end point titers, and plaque morphologies revealed a modest reduction in growth kinetics for rOSU compared to the laboratory-adapted strain. To determine if the reverse genetics system allowed for the development of the OSU virus as an expression vector, we generated rOSU with a modified segment 7 RNA that encoded the fluorescent reporter UnaG (rOSU-2A-UnaG). Immunoblot analysis and fluorescence microscopy revealed the expression of a self-cleaved and functional UnaG protein following rOSU-2A-UnaG infection. Together, this work reveals (i) the first reverse genetics system for a porcine RV, (ii) a platform for making targeted genetic modifications of OSU, and (iii) a system that allows for the expression of foreign proteins during OSU infection.

## MATERIALS AND METHODS

### Cells and virus

Embryonic monkey kidney (MA104) cells were grown in Dulbecco’s modified eagle medium (DMEM) containing 5% fetal bovine serum (FBS) and 1% penicillin-streptomycin at 37°C in a 5% CO_2_ incubator. Baby hamster kidney cells that constitutively express the T7 RNA polymerase (BHK-T7) were provided by Dr. Ulla Buchholz (Laboratory of Infectious Diseases, NIAID, NIH) and were grown in Glasgow minimum essential medium (GMEM) containing 5% heat-inactivated FBS, 1% penicillin-streptomycin, 2% nonessential amino acids, and 1% glutamine at 37°C in a 5% CO_2_ incubator. BHK-T7 cells were grown in medium supplemented with 2% Geneticin in every other passage.

RVA/Pig-tc/USA/1975/OSU/G5P[7] was provided by Dr. Taka Hoshino (Laboratory of Infectious Diseases, NIAID, NIH). This virus was activated by the addition of 10 μg/mL trypsin (final concentration) and incubating at 37°C for 60 min. The activated virus was then propagated in MA104 cells maintained in serum-free DMEM with 0.5 μg/mL porcine pancreatic trypsin (type iX, Sigma). The infected cells lysates were clarified by low-speed centrifugation at 1500×*g* for 15 min at 4°C. Virus was isolated from the clarified lysates by vertrel extraction followed by ultracentrifugation at 100,000×*g* for 2 h at 4°C. The pelleted virus [OSU-tc(MA104)] virus was resuspended in 500 μL of Tris-buffered saline and stored at −80°C.

### Nanopore sequencing

OSU-tc_(MA104) dsRNA was extracted from 250 μL of clarified infected cell lysate using a Zymo Research Direct-zol RNA Miniprep Kit following the manufacturer’s instructions. Prior to library preparation, the dsRNA was denatured with dimethylsulfoxide and poly(A)-tailed using New England Biolabs *Escherichia coli* Poly(A) polymerase following the manufacturer’s instructions. The poly(A)-tailed RNA was subjected to library preparation using an Oxford Nanopore Technologies direct cDNA sequencing kit (SQK-DCS109) following the manufacturer’s instructions. The library was sequenced using an Oxford Nanopore Technologies MinION sequencer. Sequence assemblies of the 11 genome segments of OSU-tc_(MA104) were prepared using Geneious Primer software version 2023.2.1.

### OSU sequences used in the generation of T7 expression plasmids

The sequences of the OSU-tc_(MA104) genome segments were deposited in GenBank under the accession numbers OP978238-OP978248. The sequences of modified segment 7 RNAs of OSU NSP3-2A (PP112343) and OSU NSP3-2A-UnaG (PP112344) were also deposited in GenBank.

### Plasmids used in this study

Recombinant SA11 (rSA11) viruses were prepared using the plasmids pT7/SA11VP1, pT7/SA11VP2, pT7/SA11VP3, pT7/SA11VP4, pT7/SA11VP6, pT7/SA11VP7, pT7/SA11NSP1, pT7/SA11NSP2, pT7/SA11NSP3, pT7/SA11NSP4, and pT7/SA11NSP5 and pCMV/NP868R (1, 28). Recombinant OSU (rOSU) viruses were prepared using the plasmids pT7/OSUVP1, pT7/OSUVP2, pT7/OSUVP3, pT7/OSUVP4, pT7/OSUVP6, pT7/OSUVP7, pT7/OSUNSP1, pT7/OSUNSP2, pT7/OSUNSP3, pT7/OSUNSP4, and pT7/OSUNSP5 and pCMV/NP868R (28). The plasmid pT7/SA11 NSP3-2A-UnaG was previously described (17). The plasmids pT7/OSU NSP3-2A and pT7/OSU NSP3-2A-UnaG were produced by fusing DNA fragments for 2A or 2A-3xFLAG-UnaG, respectively, to the 3’-end of the OSU NSP3 open reading frame of pT7/OSU NSP3 using the TaKaRa Bio In-Fusion cloning system. Primer synthesis and plasmid sequencing were performed by EuroFins Scientific (TABLE 1).

**TABLE 1.**
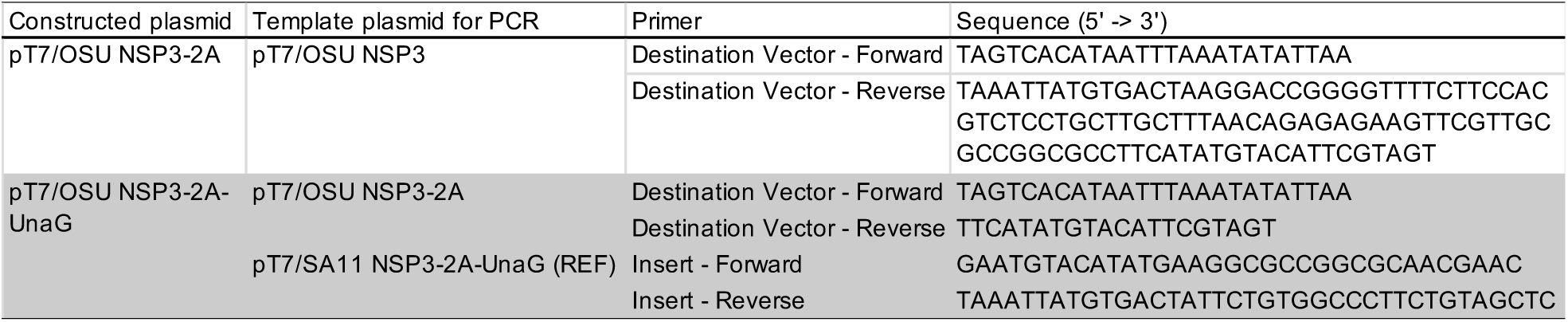
Primers used to construct pT7/OSU NSP3-2A and pT7/OSU NSP3-2A-UnaG by In-Fusion cloning.

### Isolation, amplification, and analysis of recombinant viruses

Rotavirus reverse genetics was performed as previously described (1, 29). Briefly, BHK-T7 cells in 12-well plates were transfected with the 11 SA11 or OSU T7 plasmids, or combinations thereof, and with the capping enzyme plasmid, pCMV-NP868R, using Mirus TransIT-LT1 transfection reagent. Transfection mixtures contained 0.8 μg each of the pT7 plasmids except for pT7/NSP2 and pT7/NSP5 which were used at 3-fold higher concentrations.

Two days post transfection, the BHK-T7 cells were overseeded with MA104 cells, and trypsin was added to the medium to a final concentration of 0.5 μg/mL. Three days later, the BHK-T7/MA104 cell mixtures were freeze-thawed thrice, and the lysates were clarified by low-speed centrifugation at 800×*g* for 5 min at 4°C. To amplify the recovered viruses, the lysates were adjusted to 10 μg/mL trypsin and incubated for 1 h at 37°C. MA104 cells in 6-well plates were then infected with 300 μL of the trypsin-treated lysates and incubated at 37°C in a 5% CO_2_ incubator until all cells were lysed (typically 3-5 days). Recombinant viruses were recovered from the lysates by plaque isolation on MA104 cells. Plaque-isolated viruses were initially grown on MA104 cells in 6-well plates and then, to generate larger pools, grown on MA104 cells in T175 tissue culture flasks at low multiplicity of infection (<1 PFU/cell). Briefly, 100 μL of the plaque-amplified lysates were activated by incubation with 10 μg/mL trypsin (final concentration). The activated lysates were diluted into 10 mL of serum-free DMEM and then used as inoculum to infect MA104 cells in T175 flasks. The flasks were placed at 37°C in a 5% CO_2_ incubator for 1 h with rocking to ensure equal coverage of the inoculum over the monolayers. Following adsorption, the inoculum was removed and 25 mL of serum-free DMEM containing 0.5 μg/mL trypsin was added to each flask. The flasks were returned to the incubator until all cells were lysed (typically 3-5 days). The infected cell lysates were collected, clarified by low-speed centrifugation at 800×*g* for 5 min at 4°C, and stored at −80°C. Viral dsRNAs were recovered from the infected cell lysates by TRIzol extraction, resolved by electrophoresis on 8% polyacrylamide gels in Tris-glycine buffer, detected by staining with ethidium bromide, and visualized using a Bio-Rad ChemiDoc MP imaging system. Peak titers were determined from the infected cell lysates by plaque assay.

### Plaque assay

Rotavirus plaque assays were performed as previously described (29). At 5 days post infection, MA104 monolayers with agarose overlays were incubated overnight with phosphate-buffered saline (PBS) containing 3.7% formaldehyde. The agarose overlays were then removed, and the monolayers were stained with 1% crystal violet in 5% ethanol for 3 h. The fixed and stained monolayers were rinsed with water and air dried. The plaque diameters were measured using ImageJ software (30). Statistically significant differences in titer and plaque size were determined using ANOVA (GraphPad Prism).

### Immunoblot analysis

Rotavirus infections were performed as previously described (29). Briefly, MA104 cells in 6-well plates were infected with 5 PFU/cell of the indicated viruses. At 9 h post infection, the infected cells were scraped into cold PBS. The infected cells were then washed with cold PBS, pelleted by centrifugation at 5,000×*g* for 5 min at 4°C, and lysed by incubation with nondenaturing lysis buffer (300 mM NaCl, 100 mM Tris-HCl [pH 7.4], 2% Triton X-100, and 1×EDTA-free protease inhibitor cocktail [Roche Complete]) for 30 min on ice. For immunoblot analysis, lysates were resolved by electrophoresis on 10% polyacrylamide gels in Tris-glycine buffer and transferred to nitrocellulose membranes. After blocking with PBS containing 0.1% Tween-20 and 5% nonfat dry milk, the blots were probed with FLAG M2 antibody (F1804, Sigma Aldrich, 1:2,000), 2A antibody (NBP2-59627, Novus, 1:1,000), rotavirus NSP3 antibody (lot 55068, 1:2,000), rotavirus VP6 antibody (lot 53963, 1:2,000), or β-actin antibody (D6A8, Cell Signaling Technology, 1:2000). The bound primary antibodies were detected using 1:10,000 dilutions of horseradish peroxidase (HRP)-conjugated secondary antibodies (goat anti-mouse IgG [CST], goat anti-guinea pig IgG [KPL], or goat anti-rabbit IgG [CST]) in 5% nonfat dry milk. HRP signals were developed using the Bio-Rad Clarity Western ECL substrate and developed using a Bio-Rad ChemiDoc imaging system.

### Genetic stability analysis

Genetic stability experiments were performed as previously described (3, 16–18). Briefly, the indicated viruses were serially passaged five times using 1:100 dilutions of infected cell lysates that were prepared in serum-free DMEM. When cytopathic effect reached completion (typically 3-5 days), the cells were freeze-thawed thrice. Viral dsRNAs were recovered from the infected cell lysates by TRIzol extraction (29), resolved by electrophoresis on 8% polyacrylamide gels in Tris-glycine buffer, detected by staining with ethidium bromide, and visualized using a Bio-Rad ChemiDoc MP imaging system.

### Assessment of infectivity by infectious particle production

Rotavirus infections were performed as previously described (29). Briefly, MA104 cells in 6-well plates were infected with 5 PFU/cell of the indicated viruses. Following adsorption, the infected cells were washed thrice with PBS and incubated in serum-free DMEM containing 0.5 μg/mL trypsin at 37°C in a 5% CO_2_ incubator. At the indicated times post infection, the infected cells were freeze-thawed thrice, clarified by low-speed centrifugation at 800×*g* for 5 min at 4°C, and analyzed by plaque assay. Statistically significant differences in titer were determined using ANOVA.

### Assessment of fluorescent reporter expression

Rotavirus infections were performed as previously described (29). Briefly, MA104 cells in 6-well plates were infected with 0.05 PFU/cell of rOSU and rOSU-2A-UnaG. Following adsorption, the infected cells were washed thrice with PBS and incubated in serum-free DMEM containing 0.5 μg/mL trypsin at 37°C in a Sartorius IncuCyte S3 Live-Cell Analysis System. At 28 hours post infection, images were acquired at 10X magnification under phase and green (excitation [440-480 nm], emission [504-544 nm]) channels.

### Statistical analyses

The results from all experiments represent three biological replicates. Horizontal bars indicate the means. Error bars indicate the standard deviations. *P* values were calculated using one-way analysis of variance with Bonferroni Correction (Graph Pad Prism).

## RESULTS AND DISCUSSION

### Recovery of OSU G5P[7] porcine rotavirus by reverse genetics

Genotype G5P[7] is representative of most rotaviruses (RV) that cause acute gastroenteritis in suckling and weaned pigs, which leads to economic losses that plague the global pork industry (10, 11). Current treatments are generally ineffective at preventing disease (31–33); thus, a need exists for robust molecular tools to develop next-generation porcine rotavirus vaccines. Reverse genetics systems exist for several *Rotavirus A* strains (1–9); however, no such system is available for a porcine RV. To address this knowledge gap, we utilized the well-studied Ohio State University (OSU) G5P[7] prototype strain (25–27). Laboratory-adapted OSU (OSU-tc_(MA104)) was grown in MA104 cells, and viral dsRNA was extracted from infected cell lysates. The isolated RNA was then processed for Nanopore sequencing to obtain the complete sequences of all 11 genome segments (deposited in GenBank). Notably, the Nanopore sequences for 7 OSU-tc_(MA104) genome segments (VP2, VP3, VP6, VP7, NSP2, NSP3, and NSP5) were identical to the virulent (RVA/Pig-tc/USA/1975/OSU/G5P7/virulent) and attenuated (RVA/Pig-tc/USA/1975/OSU/G5P7/attenuated) strains. In contrast, VP1, VP4, and NSP1 were >99% identical, whereas NSP4 was >95% identical (nucleotide and amino acid sequence comparisons) (27). The OSU-tc_(MA104) sequencing information was used to construct plasmids for reverse genetics experiments. Full-length cDNAs of each genome segment were positioned within T7 plasmids in between of an upstream T7 polymerase promoter and a downstream hepatitis delta virus ribozyme. In the presence of the T7 RNA polymerase, the OSU T7 plasmids produce full-length, positive sense RNA with authentic 5’ and 3’ termini.

To confirm that each OSU T7 expression plasmid was functional, we generated 11 recombinant SA11/OSU (rSA11/OSU) monoreassortants. SA11 represents the prototype strain of simian RV (34–36). Reverse genetics experiments were performed as previously described (FIG 1A) (1, 29). Briefly, 1 OSU T7 expression plasmid, 10 SA11 T7 expression plasmids, and pCMV-NP868R were transfected into BHK-T7 cells. The recovered virus was then amplified and analyzed for the production of viral dsRNA. All monoreassortants produced infectious virus (i.e., viral dsRNA) and induced cytopathic effect within 5 days of infection. Thus, the genetic information obtained by Nanopore sequencing was functional for reverse genetics. By comparing the banding patterns to rSA11, we also determined the migration distance for each rOSU genome segment (FIG 1B, see red arrows). Functional differences between the SA11 and OSU genome segments and protein products may be investigated using the monoreassortants discussed here. Strikingly, we recovered recombinant virus that was comprised of all 11 OSU genome segments (FIG 1B, see rOSU lane). Thus, we have developed a reverse genetics system for G5P[7] porcine RV that may be used for the production of vaccines targeting the most common cause of porcine RV infections. In contrast to current vaccines against porcine RV, which have been developed by serially passaging virulent strains in tissue culture - a costly and time consuming process, the OSU reverse genetics system allows for the rapid generation of vaccine candidates and for the introduction of directed attenuating genetic mutations (10, 37–40).

**FIG 1.**
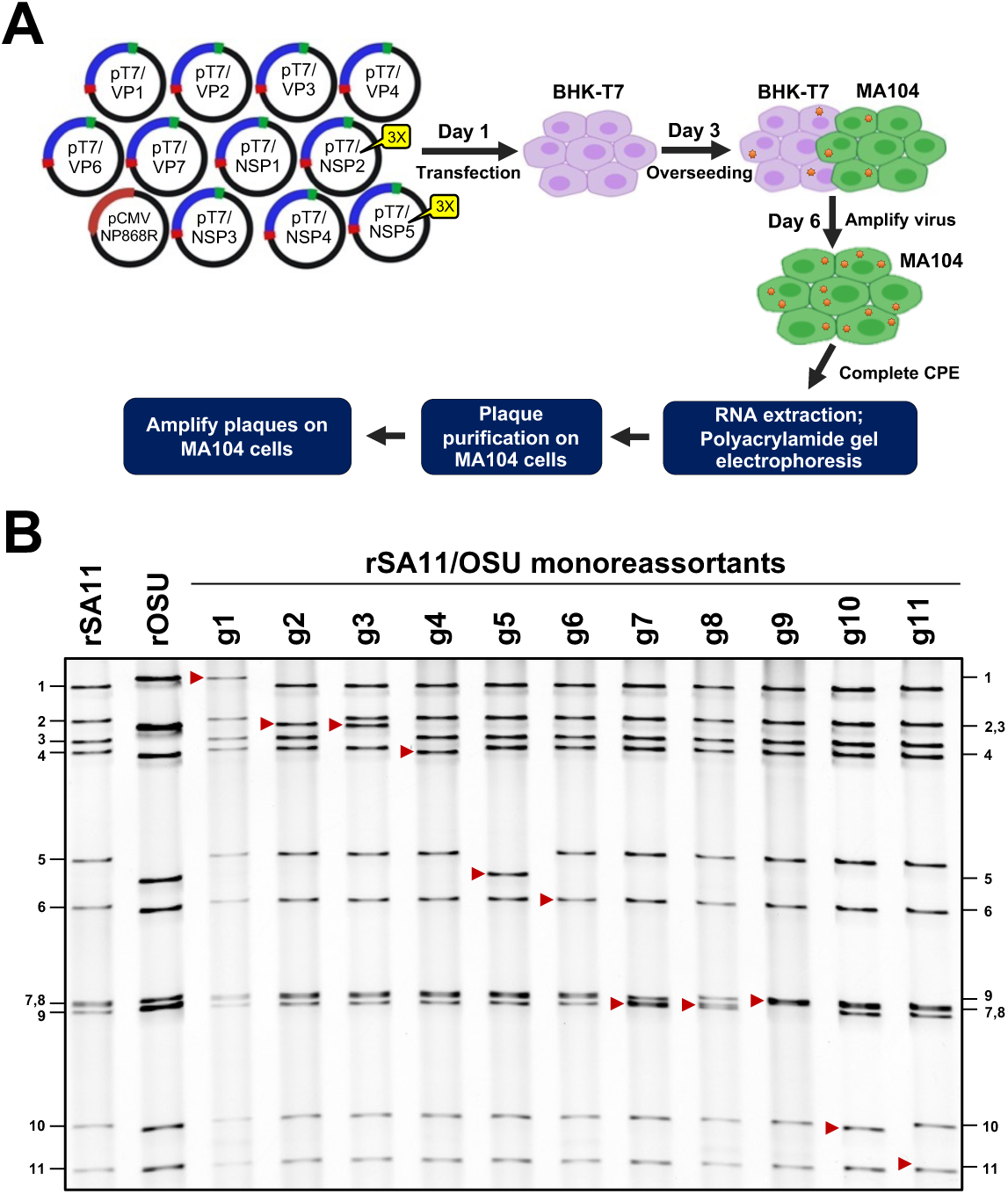
Production of OSU G5P[7] porcine rotavirus. (A) Rotavirus reverse genetics system (29). Recombinant rotavirus was prepared by transfecting BHK-T7 cells with 11 T7 plasmids, which contain full-length cDNAs of rotavirus genome segments, and the CMV-NP868R plasmid, which encodes the African Swine Fever Virus capping enzyme (28). The BHK-T7 cells were overseeded 2 days post-transfection with MA104 cells to facilitate the spread and amplification of recombinant rotavirus. At 3 days post-overseeding, recombinant virus in the cells lysates was amplified on MA104 cells. The amplified virus was analyzed by RNA gel electrophoresis followed by plaque purification. (B) Recovery of recombinant SA11/OSU monoreassortants by reverse genetics. Viral dsRNAs from rSA11, rOSU, and 11 rSA11/OSU monoreassortants were resolved by electrophoresis on an 8% polyacrylamide gel and stained with ethidium bromide. The migrations of rOSU gene segments in rSA11/OSU monoreassortants are indicated with red arrows. Genome segments 1-11 of rSA11 are indicated on the left side of the panel. Genome segments 1-11 of rOSU are indicated on the right side of the panel.

### Recovery of OSU G5P[7] porcine rotavirus encoding a foreign protein

RV can be modified to express fluorescent reporters, such as UnaG [green], mRuby [red], and TagBFP [blue], and other viral proteins, such as norovirus VP1 and SARS-COV-2 S1 (2, 3, 15–20). These recombinant viruses are valuable for analyzing RV biology by fluorescence-based imaging and may be used to induce protective immunity responses against multiple pathogenic viruses. As a proof-of-concept, we explored the possibility of expressing UnaG from OSU genome segment 7. The porcine teschovirus 2A-like (2A) element was positioned at the 3’ end of the NSP3 coding sequence followed by the in-frame coding sequence of FLAG-tagged UnaG. Due to the activity of the 2A element (22–24), translation of the RNA produces two proteins: NSP3 with remnants of the 2A element and FLAG-tagged UnaG. We generated an additional expression plasmid with only the 2A element following NSP3 (FIG 2A). Using RV reverse genetics (FIG 1A), we recovered the rOSU-2A and rOSU-2A-UnaG viruses. For each, we observed a shift in the migration pattern of genome segment 7 to a higher molecular form (FIG 2B, see red arrows). These shifts are presumably due to the extra genetic material introduced into the segment 7 RNA by addition of the 2A and 2A-FLAG-UnaG coding sequences. We also noted similar dsRNA banding patterns for OSU-tc_(MA104) and rOSU, providing additional evidence for the utility of the OSU reverse genetics system.

**FIG 2.**
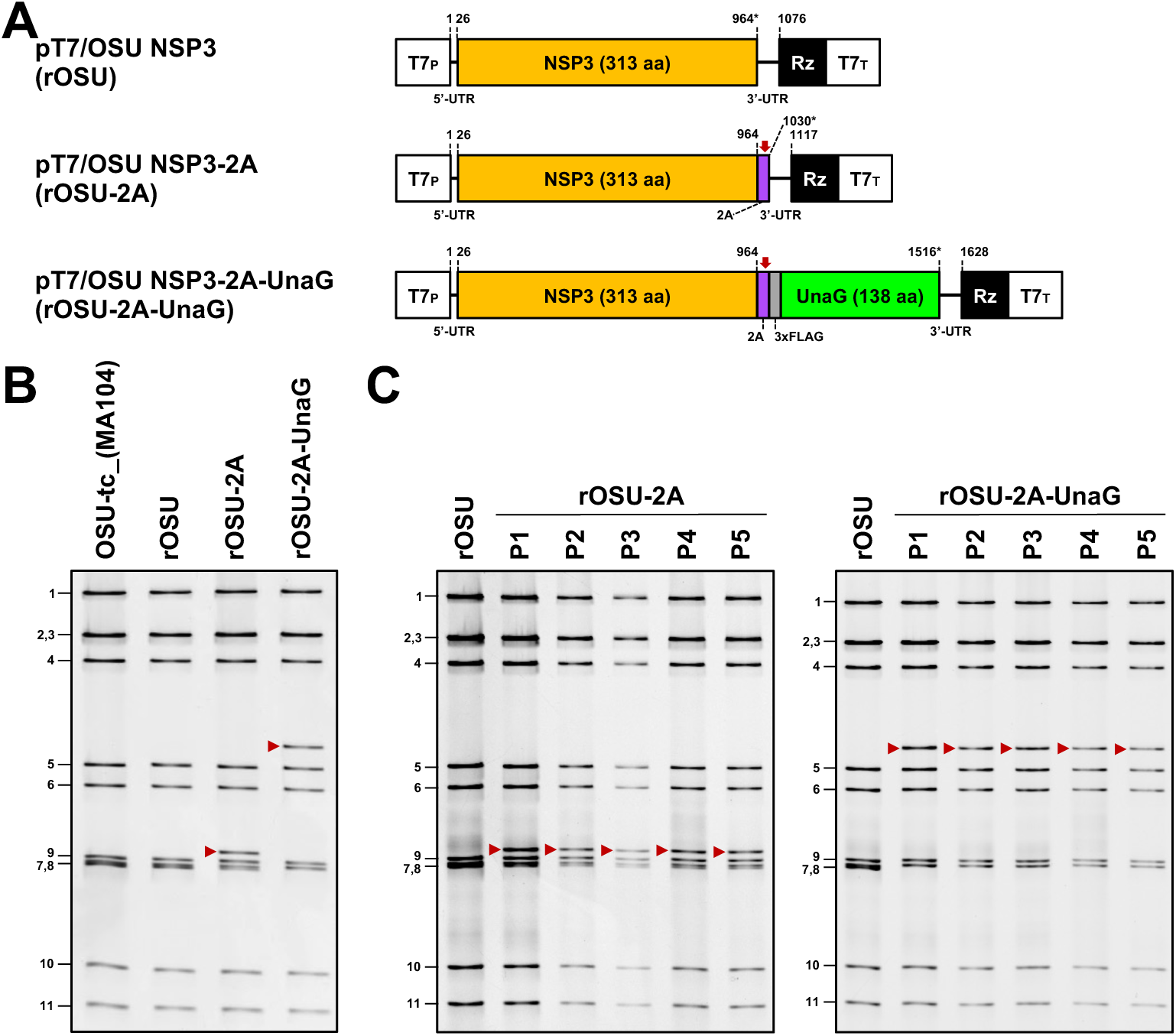
Production of recombinant OSU that encodes a foreign protein. (A) Modifications of rotavirus genome segment 7. The schematics indicate the nucleotide positions of the coding sequences for NSP3, the porcine teschovirus 2A element, 3X FLAG, and the fluorescent reporter UnaG (green). The red arrows indicate the positions of the 2A translational stop-restart elements, and the asterisks indicate the ends of the open reading frames. (B) Recovery of recombinant OSU-2A-UnaG by reverse genetics. Viral dsRNAs from OSU-tc_(MA104), rOSU, rOSU-2A, and rOSU-2A-UnaG were resolved by electrophoresis on an 8% polyacrylamide gel and stained with ethidium bromide. The migrations of modified genome segment 7 are indicated with red arrows. Genome segments 1-11 of rOSU are indicated on the left side of the panel. (B) Genetic stability. rOSU-2A and rOSU-2A-UnaG were serially passaged on MA104 cells. Viral dsRNAs from a total of 5 passages (P) were resolved by electrophoresis on an 8% polyacrylamide gel and stained with ethidium bromide. The migrations of modified genome segment 7 are indicated with red arrow. Genome segments 1-11 of rOSU are indicated on the left side of the panels.

To develop OSU G5P[7] porcine rotavirus as a vector for expressing foreign protein, the recombinant virus must be genetically stable. This property is essential for scaling large quantities for vaccine production and for other applications. To examine genetic stability, we serially passaged rOSU-2A and rOSU-2A-UnaG at low multiplicity of infection. We observed no differences in the pattern of viral dsRNA recovered from the infected lysates during passage (FIG 2C, see red arrows). The 2A-FLAG-UnaG coding sequence is stable within the OSU background for at least 5 rounds of infection; however, larger inserts may be unstable. As such, future studies will evaluate the limits of OSU genome flexibility.

### Growth characteristics of rOSU G5P[7] rotaviruses

The recombinant viruses generated in this work were derived from sequencing information of the laboratory-adapted strain. To determine if these viruses exhibit similar growth characteristics, we compared their growth kinetics and plaque morphologies. The analysis showed that rOSU was a well growing virus, reaching peak titers of ∼10^7^ in MA104 cells. However, the peak titer produced reached by rOSU was ∼0.5 log less than that reached by the OSU-tc_(MA104) virus. Moreover, single-step growth experiments indicated that rOSU grew slower than the OSU-tc_(MA104) virus. Plaque analysis also showed that the rOSU virus formed smaller plaques on MA104 cells than the OSU-tc_(MA104) virus (FIG 3A-D). These results suggest sequence differences exist between rOSU and OSU-tc_(MA104) that impact virus growth. We conclude from this that the consensus sequence information generated for OSU-tc_(MA104) genome by Nanopore sequencing may not fully reflect the distribution and combinations of sequence variations in the OSU-tc_(MA104) population associated with the fittest, best growing viruses. Indeed, the OSU-tc_(MA104) population likely represents a quasispecies, of which rOSU may, or may not, be a single variant.

**FIG 3.**
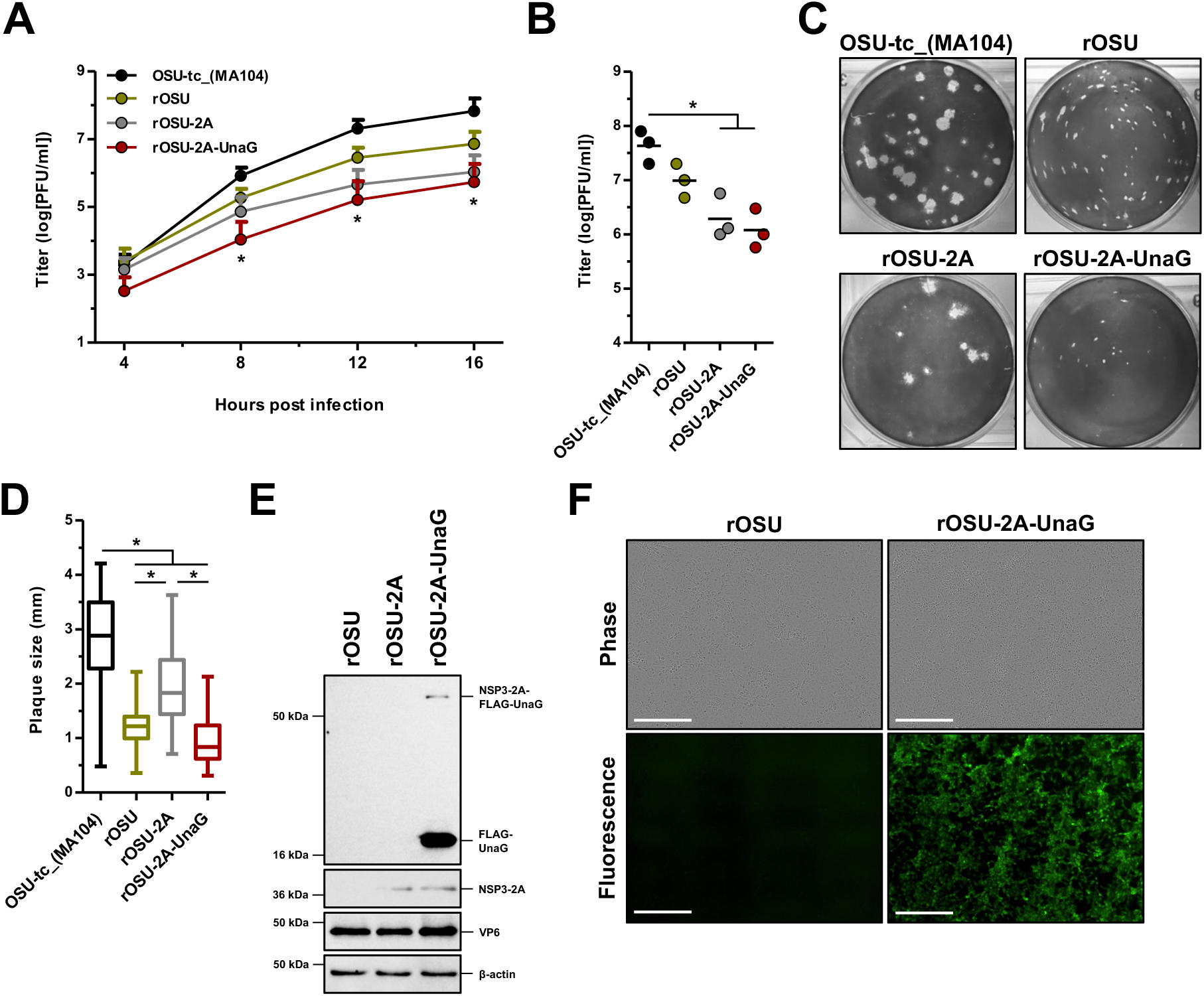
Characterization of recombinant OSU that expresses a foreign protein. (A and B) Production of infectious virus. MA104 cells were infected with the indicated viruses at an MOI of 5 PFU/cell. In panel A, titers were determined by plaque assay at the indicated times post infection. In panel B, titers were determined by plaque assay upon complete cytopathic effect (typically 3-5 days). Error bars indicate the standard deviations (A); horizontal bars indicate the means (B); in panel A, statistical analyses show the comparisons between OSU-tc_(MA104) and rOSU, rOSU-2A, and rOSU-2A-UnaG; *, P < 0.05 (n = 3 biological replicate). (C and D) Plaque morphologies and sizes. In panel C, plaques on MA104 cells were detected by crystal violet staining. In panel D, plaque diameters were measured using ImageJ software (30). 50 plaques were measured for each virus; *, P < 0.05 (n = 3 biological replicates). (E and F) Fluorescent reporter expression and activity of the 2A translational stop-restart element. In panel E, MA104 cells were infected with the indicated viruses at an MOI of 5 PFU/cell. At 9 hours post infection, infected cell lysates were prepared and analyzed by immunoblot. The migrations of molecular weight markers are indicated on the left side of the panels. The migrations of NSP3-2A-FLAG-UnaG, FLAG-UnaG, and NSP3-2A are indicated on the right side of the panels (n = 3 biological replicates). In panel F, MA104 were infected with the indicated viruses at an MOI of 0.05 PFU/cell. At 28 hours post infection, the infected cells were imaged at 10X magnification under phase and green channels. Scale bars represent 400 µm (n = 3 biological replicates).

We next examined the growth characteristics of recombinant viruses that encode foreign protein. Compared to the rOSU, rOSU-2A-UnaG produced ∼1 log-unit less infectious virus but generated similar plaque sizes (FIG 3A-D). This result is consistent with previous studies that show addition of genetic material to the segment 7 RNA is correlated with reduced vurus growth (3, 16–18).

### Expression of foreign protein by rOSU G5P[7] porcine rotavirus

To develop OSU as a dual vaccine platform, the vector must express a foreign protein. As such, the protein products made by rOSU-2A-UnaG following infection were probed by immunoblot with an anti-FLAG antibody (FIG 3E). This assay revealed high levels of cleaved FLAG-UnaG (∼18 kDa); rOSU-2A-UnaG directed expression of the foreign protein and a functional 2A element. We also detected minor amounts of uncleaved NSP3-2A-UnaG (∼56 kDa), which likely represents a readthrough of the 2A element. Probing with an anti-2A antibody revealed a protein product that migrated at the expected molecular weight for NSP3 linked to the remnant residues of the 2A peptide (∼38 kDa) (FIG 3E). Viral protein VP6 and host protein β-actin were detected under all infection conditions; however, NSP3-2A-UnaG, FLAG-UnaG, and NSP3-2A were not present in rOSU infected cell lysates. Finally, we used fluorescence microscopy to determine if the expressed and cleaved FLAG-UnaG was functional. In contrast to rOSU, rOSU-2A-UnaG produced high levels of fluorescent signal within live cells (FIG 3F).

### Conclusions

The establishment of more effective porcine RV vaccines can be advanced through the application of reverse genetics technologies to create candidates that are superior in generating protective immunological responses. Such reverse genetics technologies allow for targeted genetic modifications and avoid the costly and time-consuming process of attenuating virulent strains through serially passage. Moreover, it may be possible to design combination vaccines that are capable of protecting individuals against RV and other pathogenic viruses. In this work, we developed a robust reverse genetics system for the G5P[7] OSU strain of porcine rotavirus. Using information obtained by Nanopore sequencing the laboratory-adapted strain, we constructed 11 T7 expression plasmids that were sufficient for generating recombinant OSU (FIG 1). We leveraged the reverse genetics system to produce a recombinant virus that expressed UnaG within infected cells (FIGS 2 and 3). Recombinant viruses that express fluorescent reporters may be used as diagnostic tools for detecting infection and for measuring immunological responses. Overall, this work describes the first reverse genetics system for a porcine rotavirus and advances the development of next-generation vaccines.

## ACKNOWLEDGEMENTS

We thank members of the Indiana University virology community for helpful comments and suggestions. A special thanks to Dr. Ulla Buchholz (Laboratory of Infectious Diseases, NIH-NIAID) for providing the BHK-T7 cells and Dr. Taka Hoshino (Laboratory of Infectious Diseases, NIH-NIAID) for providing RVA/Pig-tc/USA/1975/OSU/G5P[7].

## CONFLICTS OF INTEREST

AJS, CAA, and JTP are inventors of an Indiana University patent application related to the content of this work. JTP has an interest in biotechnology companies developing vaccines using recombinant rotaviruses.

## AUTHOR CONTRIBUTIONS

Conceived and designed the experiments: AJS, CAA, and JTP. Performed the experiments: AJS and CAA. Analyzed the data: AJS, CAA, and JTP. Wrote the paper: AJS, CAA, and JTP.

## Notes

### Competing Interest Statement

The authors have declared no competing interest.

